# The *Candida auris* Hog1 MAP kinase is essential for the colonization of murine skin and intradermal persistence

**DOI:** 10.1101/2024.03.18.585572

**Authors:** Raju Shivarathri, Manju Chauhan, Abhishek Datta, Diprasom Das, Adela Karuli, Sabrina Jenull, Karl Kuchler, Shankar Thangamani, Anuradha Chowdhary, Jigar V. Desai, Neeraj Chauhan

**Affiliations:** Center for Discovery and Innovation, Hackensack Meridian Health, 111 Ideation Way Nutley New Jersey 07110, USA; Medical University Vienna, Department of Medical Biochemistry, Max Perutz Labs Vienna, Campus Vienna Biocenter, Dr. Bohr-Gasse 9/2; A-1030 - Vienna, Austria; Department of Medical Mycology, Vallabhbhai Patel Chest Institute, University of Delhi, Delhi, India; Department of Comparative Pathobiology, Purdue University College of Veterinary Medicine, 625 Harrison Street West Lafayette, Indiana 47906

**Author notes:** To whom correspondence should be addressed: Anuradha Chowdhary;, Jigar Desai;, Neeraj Chauhan.

**Keywords:** *Candida auris*, HOG1, MAP kinase, skin colonization, intradermal infection, biofilm

## Abstract

*Candida auris*, a multidrug-resistant human fungal pathogen, was first identified in 2009 in Japan. Since then, systemic *C. auris* infections have now been reported in more than 50 countries, with mortality rates of 30-60%. A major contributing factor to its high inter- and intrahospital clonal transmission is that *C. auris,* unlike most *Candida* species, displays unique skin tropism and can stay on human skin for a prolonged period. However, the molecular mechanisms responsible for *C. auris* skin colonization, intradermal persistence, and systemic virulence are poorly understood. Here, we report that *C. auris* Hog1 mitogen-activated protein kinase (MAPK) is essential for efficient skin colonization, intradermal persistence, as well as systemic virulence. RNA-seq analysis of wildtype parental and *hog1*Δ mutant strains revealed marked down-regulation of genes involved in processes such as cell adhesion, cell-wall rearrangement, and pathogenesis in *hog1*Δ mutant compared to the wildtype parent. Consistent with these data, we found a prominent role for Hog1 in maintaining cell-wall architecture, as the *hog1*Δ mutant demonstrated a significant increase in cell-surface β-glucan exposure and a concomitant reduction in chitin content. Additionally, we observed that Hog1 was required for biofilm formation *in vitro* and fungal survival when challenged with primary murine macrophages and neutrophils *ex vivo*. Collectively, these findings have important implications for understanding the *C. auris* skin adherence mechanisms and penetration of skin epithelial layers preceding bloodstream infections.

**Importance:** *Candida auris* is a World Health Organization (WHO) fungal priority pathogen and an urgent public health threat recognized by the Centers for Disease Control and Prevention (CDC). *C. auris* has a unique ability to colonize human skin. It also persists on abiotic surfaces in healthcare environments for an extended period of time. These attributes facilitate the inter- and intrahospital clonal transmission of *C. auris*. Therefore, understanding *C. auris* skin colonization mechanisms are critical for infection control, especially in hospitals and nursing homes. However, despite its profound clinical relevance, the molecular and genetic basis of *C. auris* skin colonization mechanisms are poorly understood. Herein, we present data on the identification of the Hog1 MAP kinase as a key regulator of *C. auris* skin colonization. These findings lay foundation for further characterization of unique mechanisms that promote fungal persistence on human skin.

## Introduction

Invasive fungal diseases are responsible for ∼2.5 million deaths per year worldwide – a number that far exceeds the deaths caused either by tuberculosis or malaria (1). This number is expected to rise due to the increasing number of immunosuppressed people, including the elderly, premature infants, transplant recipients, and cancer and HIV/AIDS patients (1). The Centers for Disease Control and Prevention (CDC) and the World Health Organization (WHO) have both recognized *C. auris* as an urgent threat to human health and have also recently emphasized a critical need for the development of new antifungal therapeutics to address the growing issue of antifungal drug resistance among human fungal pathogens (2–5).

*Candida* species are the foremost clinically relevant invasive fungal pathogen in the United States and a leading cause of bloodstream infections worldwide (1, 6). The genus *Candida* contains more than 30 species that can cause life-threatening human infections (7, 8). *Candida* species, unlike other major fungal pathogens, are commensal colonizers and a normal component of the human microflora present on mucosal and epithelial barriers such as the gastrointestinal and urogenital tracts (7, 9).

*Candida auris* is a newly emerging multidrug-resistant fungal pathogen first reported in Japan in 2009 (10–12). Within a decade, *C. auris* has spread around the globe, causing widespread hospital outbreaks of candidemia in healthcare settings (13, 14). The CDC has classified *C. auris* as an urgent threat to human health due to its clinical and economic impact, high transmissibility, lack of effective antifungal treatments, and future projections of new infections over the next 10 years (2, 15). Furthermore, the WHO Global Antimicrobial Resistance Surveillance System (GLASS) has also highlighted the need for a stronger global surveillance to identify the emergence of drug resistance in *Candida* infections (16). The spread of *C. auris* is facilitated through its easy transmission by skin-to-skin contact, especially in hospital environments (17). Recent reports indicate transmission of pan- and echinocandin-resistant *C. auris* strains from healthcare facilities in Washington DC and Texas (18), suggesting *C. auris* can spread quite easily among susceptible patient populations. However, despite its clear clinical relevance, our understanding of molecular mechanisms of *C. auris* skin colonization, systemic virulence, and its impacts on the interactions with the host immune system remain poorly understood.

The fungal cell wall and its components are of interest due to its significance as a potential target for antifungal therapy and its role in host-pathogen interactions. Adhesins are proteins expressed on the fungal cell surface and are critical for the fungal cell-cell interactions (e.g. flocculation), attachment to abiotic substrates as well as interactions with the host immune cells (19). Furthermore, the majority of adhesin genes are located in the sub-telomeric regions of chromosomes where they often undergo recombination events to generate genotypic diversity, especially in the clinical isolates (19–21). This suggests that associated changes in adhesin expression may contribute to host-pathogen interactions during fungal pathogenesis. For example, members of the *TLO* gene family in *C. albicans* and epithelial adhesin (*EPA*) gene family in *C. glabrata*, display strain dependent gene size reductions or expansions (19, 20, 22). While *C. albicans* contains hyphal wall protein (*HWP1*), enhanced adhesion to polystyrene (*EAP1*) and a family of agglutinin-like sequences (*ALS*) adhesin genes (19), adhesin genes of *C. auris* and their role in host-pathogen interactions remain understudied. However, orthologs of *C. albicans* adhesin genes have been identified in *C. auris* (21). Recently a *C. auris* specific cell wall adhesin Scf1 was identified and was shown to be important for biofilm formation, skin colonization and virulence in an immunosuppressed murine model of systemic infection (23). Additionally, altered *C. auris* adhesin expression is implicated in biofilm formation (24), suggesting that *C. auris* cell wall adhesins may play a prominent role in virulence.

Hog1 is a mitogen activated protein kinase (MAPK) of the high osmolarity glycerol response (HOG) signaling pathway (25–28). The HOG pathway is activated by an upstream two-component response regulator (29, 30). For example, the Ssk1 response regulator of *Candida albicans* activates the Hog1 MAPK pathway during oxidative stress (31). The activated MAP kinase cascade, in turn, activates downstream transcription factors of genes associated with morphogenesis, adhesion, stress response, drug resistance and virulence factor expression (30). In *C. albicans* Hog1 MAPK regulates glycerol accumulation and adaptation to high osmolarity, oxidative stress, morphogenesis, and cell wall biosynthesis (32–36). Published studies from our group, as well as other investigators, have shown that the Hog1 MAPK pathway is important for stress tolerance, drug resistance and virulence of *C. albicans, Cryptococcus neoformans, Aspergillus fumigatus,* and *C. auris* (37–41). While orthologs of Hog1 have been studied in many fungal pathogens, it is important to note that there are differences in the function of Hog1 across several different fungal species studied thus far. For example, *C. albicans* Hog1 is a major regulator of oxidative stress response, a function not observed with *S. cerevisiae* Hog1 (31). Here, we show that *C. auris* Hog1 is essential for skin colonization, intradermal persistence, and systemic virulence. Interestingly, the Hog1-regulated fungal processes appear to be dispensable for driving early phagocyte recruitment to the infected tissues. In contrast, Hog1 promoted adherence and β-glucan masking, which appeared to have critical functions in promoting fungal persistence and intracellular survival within the phagocytes. Overall, our work establishes a foundation for a detailed molecular and genetic understanding of the Hog1 MAPK network in *C. auris* and its role in skin colonization and invasive infection.

## Results

### RNA-seq of *hog1*Δ reveals significant enrichment of cell wall biogenesis, adhesion, and host defense-related genes

Previously, we showed that *C. auris* Hog1 is required for antifungal drug resistance (41). To determine the Hog1-dependent transcriptional program, we performed RNA-seq analysis of logarithmically growing *hog1*Δ and wildtype parent (1184/P/15) in the Yeast Extract Peptone Dextrose broth medium at 30°C. Differentially expressed genes (DEGs) were defined by a 1.5- fold change (log2FC 0.585), with an adjusted *P* value cutoff ≤ 0.05. The complete list of differentially expressed genes in the *hog1*Δ mutant strain is presented in Tables S1. A multivariate principal-component analysis (PCA) revealed that the biological replicates from each strain clustered together, whereas the *hog1*Δ and 1184/P/15 clustered separately by principal component 1 (PC1, 44% of explained variance) (Fig. 1A). A majority of *C. auris* genes are uncharacterized. Therefore, their corresponding *C. albicans* orthologs are also not well defined and there is discrepancy in the nomenclature of most *C. auris* orthologs in the Candida Genome Database (42, 43). Where confirmed, we have provided the *C. albicans* ortholog alongside the *C. auris* gene names. A total of 2060 DEGs were found in *hog1*Δ strain compared to the wildtype 1184/P/15 strain, while no detectable reads were observed for B9J08_004369 (*HOG1*) (Fig. 1B-C). Among the DEGs, 966 were upregulated and 1094 were down regulated (Fig. 1C). Importantly, gene ontology (GO) term analysis of DEGs revealed *C. auris* genes associated with cell wall organization, adhesion, signal transduction and pathogenesis were significantly affected in *hog1*Δ compared to the wildtype parent (Fig. 1D-E). For example, cell wall genes such as B9J08_001418 (*BGL2*), B9J08_001242 (*PGA1*), B9J08_005245 (*PGA4*), B9J08_000918 (*PHR1*), B9J08_003910 (*PIR1*), and B9J08_003251 (*XOG1*) were significantly downregulated in the *hog1*Δ compared tothe wildtype (Fig. 1D). Of these cell wall genes, B9J08_001418 (*BGL2)* and B9J08_003251 (*XOG1)* are putative Δ-1,3 glucanase (44). Their primary function is to reduce the exposure of β-glucan to phagocytic cells (44). Moreover, the expression of a putative chitin synthase B9J08_003879 (*CHS2*) was also repressed in *hog1*Δ. On the other hand, the cell-wall mannosylation genes, including the secretory pathway P-type Ca^2+^/Mn^2+^-ATPase (PMR1), protein mannosyltransferase (*PMT1*), and a member of Mnn9 family mannosyltransferase (*VAN1*), were not differentially expressed (Fig. S1A). Additionally, a number of genes associated with potential cell wall functions, which are uncharacterized and lack known orthologs in other *Candida* species, were also found to be significantly differentially expressed in *hog1*Δ. A few examples of these genes include B9J08_003910, B9J08_005431, and B9J08_004476. Interestingly, among the genes with putative functions in adhesion, there was no significant change in the transcript of *SCF1* (B9J08_001458) in the *hog1*Δ mutant strain (Fig. S1B). However, the expression of B9J08_004109 (*IFF4109*) that belongs to the *IFF*/*HYR* family of adhesins was ∼2 fold downregulated. Furthermore, among the ALS (Agglutinin-Like Sequence) adhesins the expression of B9J08_002582 (*ALS4*) and B9J08_004498 (similar to *C. albicans ALS3*) were not affected while B9J08_004112 (*ALS5*) was upregulated by ∼2 fold in the *hog1*Δ compared to the wildtype. We also noted the expression of several genes associated with host recognition and/or pathogenesis affected in the *hog1*Δ mutant. The molecular chaperone *HSP90* a key regulator of fungal morphogenesis and drug resistance (45) was slightly upregulated in *hog1*Δ (Table S1). Among the most strongly downregulated genes were B9J08_004108 (putative ATPase), B9J08_001360 (putative vacuolar ATPase), B9J08_004175 (putative lipase), B9J08_001381 (*SOD1*), B9J08_002431 (*VPS28*), B9J08_003140 (*VPS51*). With the exception of *HSP90*, all of these genes are uncharacterized in *C. auris* but based on the function of their orthologs in *C*. *albicans* these genes are predicted to be important for ATPase, lipase & superoxide dismutase activity as well as vacuole transport related function (46–49). Importantly, the transcriptome data correlates well with our observation that *C. auris* Hog1 is an important regulator of adhesion, cell wall architecture and virulence.

**Fig. 1:**
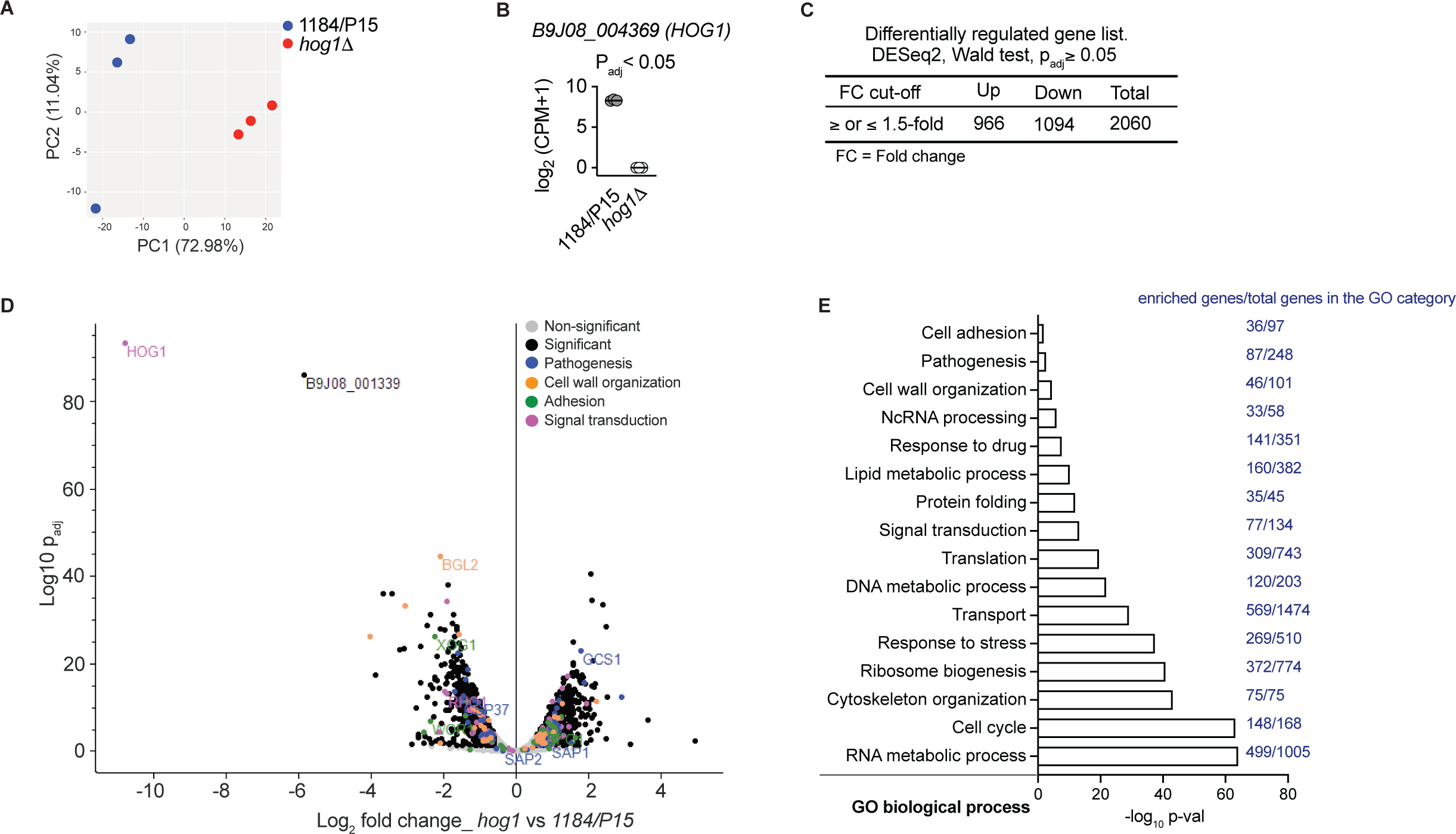
Transcriptional response of *C. auris* clinical isolate 1184/P/15 and *hog1*Δ cells. Gene expression profiles of *C. auris* isolates grown in YPD were determined using RNA-seq of three biological replicates. **(A)** Principal-component analysis (PCA) of normalized read counts from three biological replicates per strain displays the level of correlation and the reproducibility among different biological replicates. **(B)** Log-transformed normalized counts depict the variation of *HOG1* expression in parent and *hog1*Δ cells. **(C)** The number of differentially expressed genes along with up (≥1.5-fold) and downregulated (≤ −1.5-fold) genes (padj ≤ 0.05) are tabulated. Fold change 1.5 = Log_2_ FC 0.585. **(D)** Volcano plot displays the log_2_-fold change versus statistical significance (Log_10_ padj) in *hog1*Δ cells relative to the parent isolate. A total 2060 significantly enriched (≥1.5-fold, log_2_-FC ≥0.585) and de-enriched (≤-1.5-fold, log_2_-FC ≤-0.585) genes are indicated in black dots. The colored and labelled dots represent genes involved in various biological processes such as pathogenesis (blue), cell wall organization (orange), adhesion (green), and signal transduction (purple). **(E)** The significantly enriched (*p* ≤ 0.05) Gene Ontology (GO) categories (biological process) in *hog1*Δ cells compared to parent cells are illustrated. The number of genes/GO category were indicated next to GO term.

### Deletion of *HOG1* results in defective biofilm formation by *C. auris*

Based on the observed changes in the transcriptome of *hog1*Δ compared to the wildtype parental strain, and a significant down-regulation of genes belonging to the biological processes of adhesion and cell wall organization, we hypothesized that the Hog1 MAPK regulates *C. auris* adherence and biofim development. To test this hypothesis, we tested the *hog1*Δ and wildtype strains for biofilm formation on borosilicate cover glass. The *hog1*Δ formed weak biofilms, with multicellular cell clusters appearing dispersed and a discontinuous growth pattern was visible across the cover glass surface (Fig. 2A). Conversely, the wildtype strain formed robust biofilms evident from the thick uniform growth on the cover glass (Fig. 2A). The biofilm thickness was significantly reduced for *hog1*Δ. These data demonstrate that Hog1 MAPK has roles in *C. auris* adherence and efficient biofilm development.

**Fig. 2:**
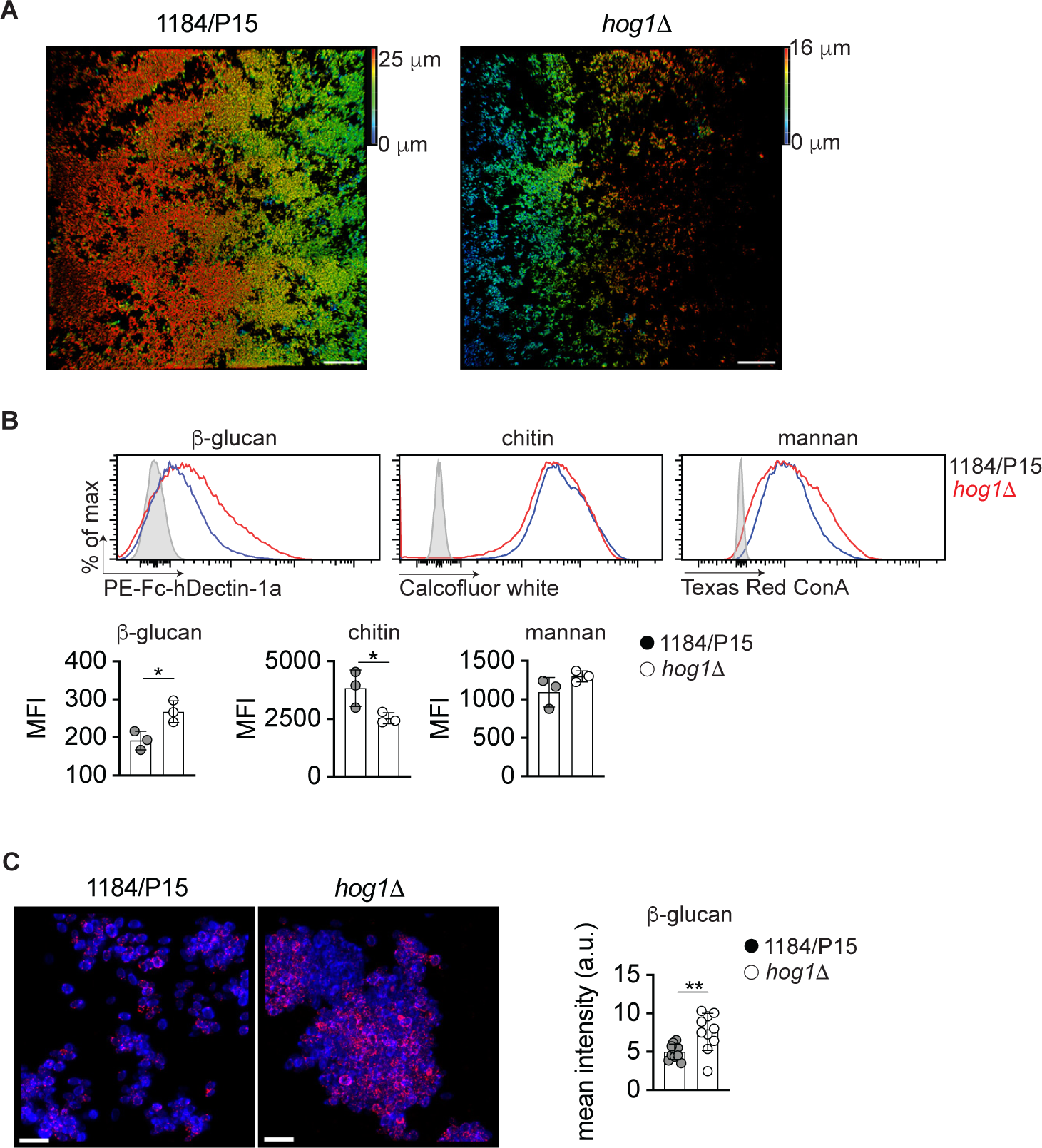
Lack of *C. auris HOG1* results in defective biofilm formation and altered cell wall composition. (A) Representative maximum intensity projection image of confocal image displaying the biofilm thickness in parent and *hog1*Δ cells. Scale bar (100 μm). **(B)** *C. auris* isolates grown in synthetic sweat media at 30°C were washed and triple-stained to quantify cell wall components. Representative flow-cytometry histograms showing β-glucan, chitin, and mannan. The quantification data are shown below the histograms form three biological replicates. Dots represent an individual replicate. Error bars represent mean ± SD; \**P* < 0.05, by unpaired *t* test. **(C)** Representative maximum intensity projection image of confocal image stacks depicting PE- FchDectin-1a stained β-glucan in RED, while calcofluor white stained chitin in BLUE. Scale bar (25 μm). Quantification of exposed β-glucan (Right). Data represent the mean fluorescence intensity. Each dot represents the mean of a single randomly chosen region of interest. Error bar represents mean ± SD; \*\**P* < 0.01, by unpaired *t* test.

### Hog1 is required for masking the *C. auris* cell wall, β-glucan

The fungal cell wall is the first point of contact with the host (19, 50). It holds an array of pathogen-associated molecular patterns (PAMPs) engaging with host pattern recognition receptors (PRRs) that mediate pathogen recognition and antifungal effector mechanisms such as phagocytosis and the release of cytotoxic reactive oxygen species (ROS) (51). Due to the observed role of Hog1 in the transcriptional regulation of genes involved in cell-wall organization (Fig. 1), we hypothesized that Hog1 might regulate the cell wall associated PAMPs. To this end, we quantified the major carbohydrate components of the fungal cell wall such as β-glucan, mannan, and chitin by using a flow cytometry-based approach (41, 52). Since *C. auris* exhibits a predilection for the dermal association, we grew the fungus in the synthetic sweat medium (53), and quantified the aforementioned cell surface carbohydrates. Our results revealed that β-glucans exposure was significantly elevated in the *hog1*Δ (Fig. 2B). To further delineate the changes in β-glucan exposure observed in the *hog1*Δ mutant, we performed fluorescence microscopy to visualize the exposed β-glucan on the cell surface in wildtype and *hog1*Δ strains (Fig. 2C). The fluorescence microscopy revealed uniformly enhanced binding of β-glucan by Fc-hDectin-1a around the cell surface in *hog1*Δ mutant strain, while the wildtype showed a discontinuous/non-uniform punctate pattern of binding with Fc-hDectin-1a (Fig. 2C). Conversely, the chitin content was reduced in *hog1*Δ compared to the wildtype parent (Fig. 2B), while no changes were observed in mannan content (Fig. 2B). Collectively, these data reveal that *C. auris* Hog1 controls the β-glucan exposure on the fungal cell surface.

### The *C. auris* Hog1 is required for intracellular survival in murine macrophages and neutrophils

Myeloid phagocytes, including macrophages and neutrophils, are key components of the innate immune system as they engage in anti-fungal host protection via the deployment of oxidative and non-oxidative mechanisms of fungal killing (54, 55). To determine whether Hog1 regulates the mechanisms to counteract the phagocytic killing, we infected murine bone marrow-derived macrophages and neutrophils with wildtype and *hog1*Δ strains at a multiplicity of infection (MOI) of 5. Surviving fungal cells were quantified by enumerating the colony-forming units (CFU) after 2.5 h of co-culture. Strikingly, *hog1*Δ was much more sensitive to killing by macrophages and neutrophils when compared to wildtype cells (Fig. 3A). Furthermore, the enhanced killing of *hog1*Δ correlated well with increased reactive oxygen species (ROS) production by macrophages and neutrophils (Fig. 3B-E) when the phagocytes were challenged with *hog1*Δ. Thus, the loss of Hog1 promoted enhanced ROS production and concomitantly reduced fungal viability inside macrophages and neutrophils.

**Fig. 3:**
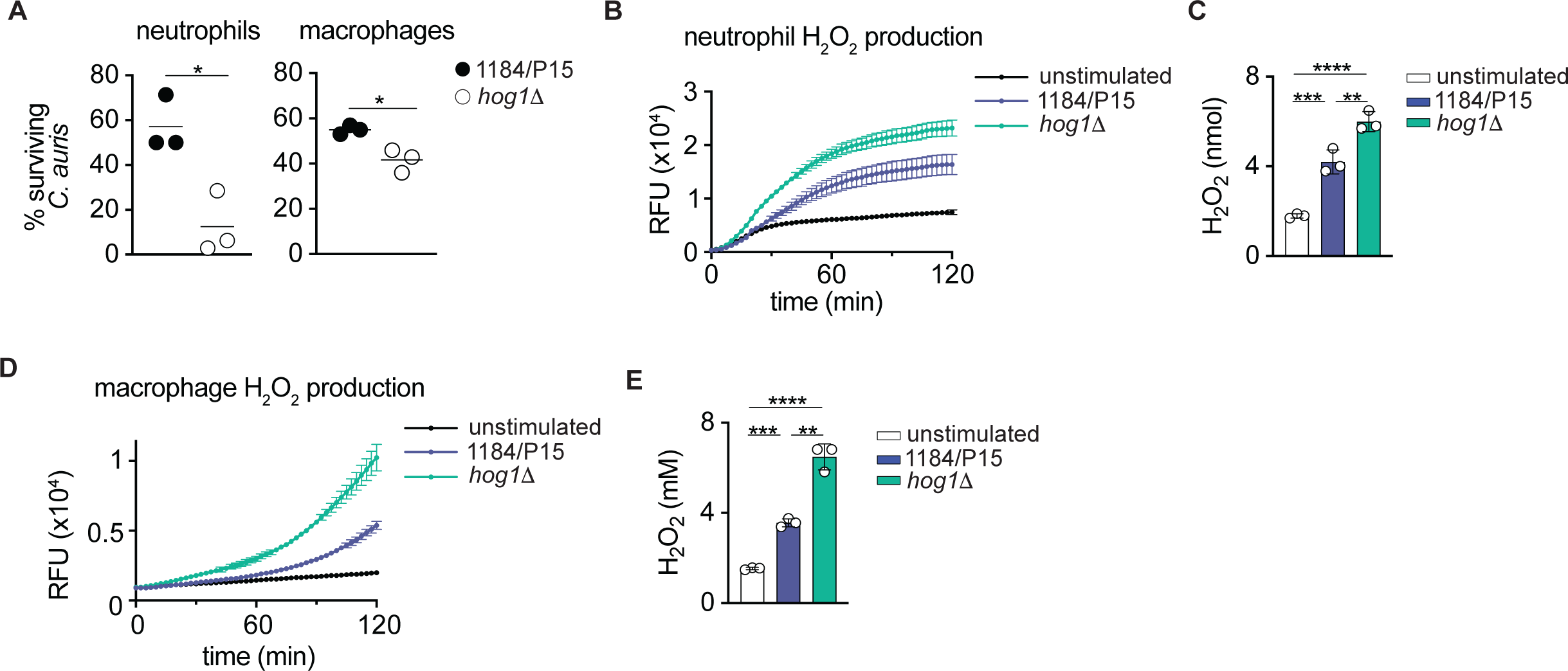
The *C. auris HOG1* is required for the survival inside bone marrow-derived macrophages and neutrophils. (A) Survival of *Candida auris* post-challenge with bone marrow-derived neutrophils or macrophages. Data represent the mean of three biological replicates (horizontal line). Dots represent an individual replicate. * p<0.05 with unpaired two-tailed *t* test. **(B-E)** Hydrogen peroxide production by neutrophils and macrophages upon the challenge with indicated *C. auris* strains. Temporal trace of hydrogen peroxide production of neutrophils (B), and macrophages (D), while the amount of the total peroxide produced after 2h of stimulation is depicted in "**C"** for neutrophils, and in "**E"** for macrophages. Data represent the mean± SD from three biological replicates, whilethe dots represent individual replicate **(C, E)** ** p<0.005, *** p<0.0005, ****p<0.0001, by one-way ANOVA followed by Tukey’s multiple comparison test **(C, E)**.

### *C. auris* Hog1 MAPK is essential for colonization of murine skin, intradermal persistence, and systemic infection

Deletion of *HOG1* results in increased β-glucan exposure, and enhanced killing by phagocytes (Fig. 2-3). Both these processes have been shown to affect the virulence of *Candida* spp (44, 56, 57). A previous report also indicated that Hog1 is required for *C. auris* virulence in a *Caenorhabditis elegans* model of infection (40). Thus, we reasoned that Hog1 impacts *C. auris* virulence *in vivo*. To determine the role of Hog1 in skin colonization, intradermal persistence, and systemic virulence, we infected immunocompetent C57BL/6 mice with *hog1*Δ and wildtype parental strains using the epicutaneous, intradermal, and systemic infection models (58–60). For skin infections (epicutaneous as well as intradermal), an equal number (6 each) of immunocompetent C57BL/6 mice were infected with 5x10^7^ *C. auris* strains via topical application on skin or with 1x10^7^ *C. auris* strains via intradermal injection. For invasive infections, an equal number (6 each) of immunocompetent C57BL/6 mice were systemically infected with 5x10^7^ *C. auris* blastospores. The animals were monitored daily for clinical signs of disease and mortality. Interestingly, there was a significant difference in fungal burdens 72 hours post-infection from mice infected with wildtype parent and *hog1*Δ strains via the epicutaneous or intradermal route (Fig. 4A-B). Moreover, histopathology of infected murine skin tissues confirms differences in the fungal burden between the wildtype and *hog1*Δ. The periodic acid-Schiff (PAS) stained skin section shows a larger fungal presence in wild type infected mice compared to the *hog1*Δ mutant (Fig. 4C-D). Similarly, mice infected with the *hog1*Δ showed significantly reduced fungal burden in the kidneys and brain, while splenic and hepatic burdens were similar to the wildtype strain (Fig. S2). Interestingly, the enhanced tissue clearance of the *hog1*Δ was not dependent upon the tissue accumulation of the myeloid phagocytes, including the neutrophils, monocytes, and macrophages, as the *hog1*Δ infected mice did not exhibit substantially different accumulation of the aforementioned myeloid phagocytes in the skin (Fig. S4B-C), same as in kidneys upon systemic infection (Fig. S4D). Hence, our data suggest that the Hog1-dependent fungal responses are necessary for countering the phagocytes’ effector functions, but do not induce tissue recruitment. Overall, these data demonstrate that Hog1 is essential for efficient epidermal colonization, intradermal persistence, and systemic infection.

**Fig. 4:**
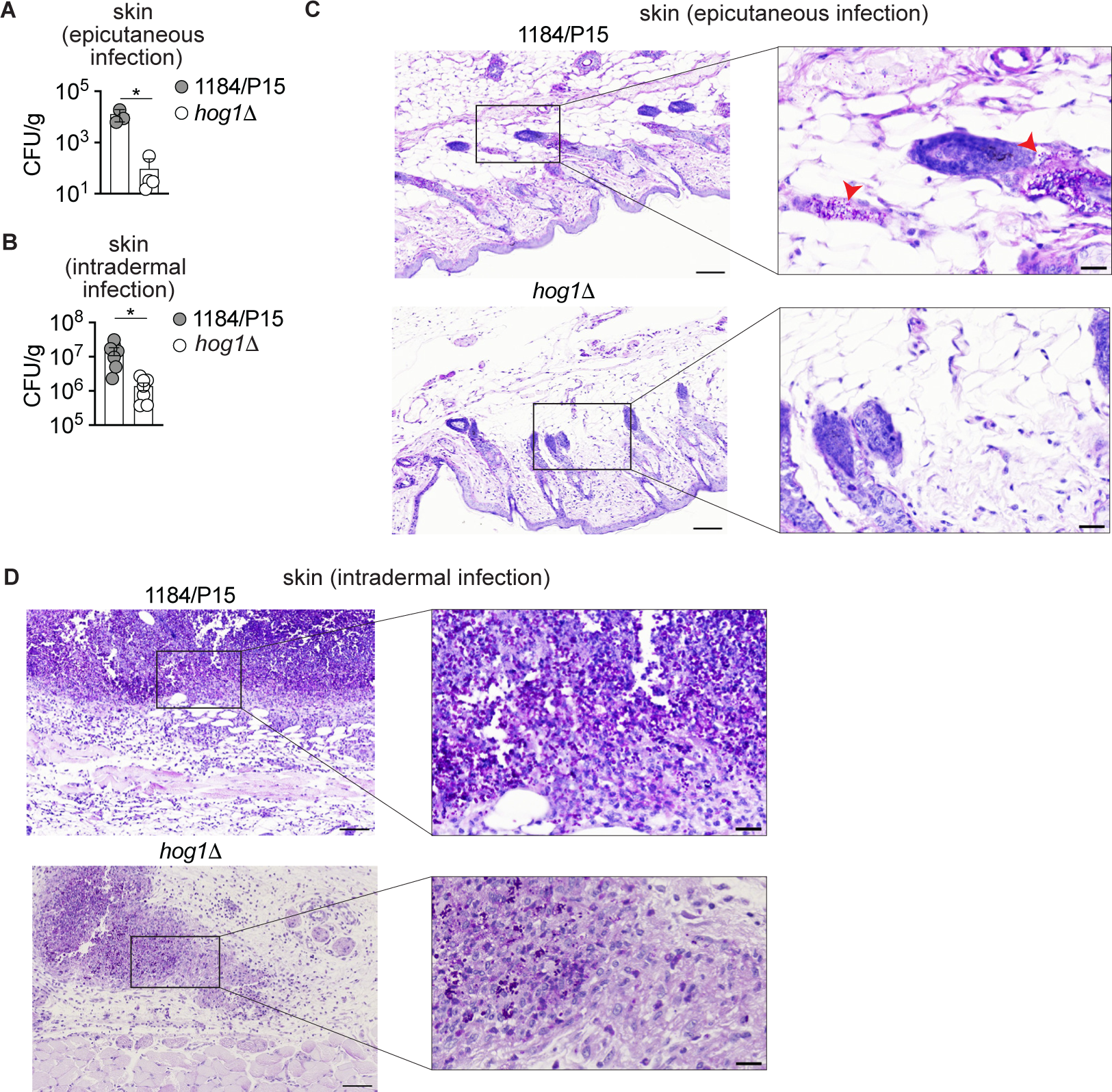
Genetic ablation of *C. auris* Hog1 MAPK impairs murine skin colonization and intradermal persistence. The C57BL/6 mouse skin was colonized with *C. auris* parent and *hog1*Δ cells epicutaneously (A and C), and intradermally (B and D) with 5x 10^7^ and 1x10^7^ CFUs, respectively. Three days post infection, animals were euthanized and collected both left-side skin (for CFUs enumeration) and right-side skin (for histopathology). (A-B) Tissue fungal burden at 72h post-infection. Data represent the mean of 4-7 individual mice from two independent experiments. Dots represent individual mice. Error bars represent mean ± SEM; \**P* < 0.05, by unpaired Mann-Whitney *U* test (A) and unpaired Welch’s *t* test (B). (C-D) Representative histopathology of skin sections stained with periodic acid-Schiff stain. Scale bar, 100 μm. Inset shows high-magnification image (scale bar 25 μm). The red arrowheads in “C” indicate *C. auris*.

## Discussion

In this study, we investigated the role of Hog1 MAPK in skin colonization and systemic infection by integrating transcriptomics, *C. auris*-host immune effector function analysis, and murine models of skin colonization, intradermal persistence, and systemic infections. Our data demonstrate that the genetic removal of *HOG1* impairs the ability of *C. aruis* to colonize and persist on murine skin as well as cause successful systemic infections. Furthermore, the deletion of *HOG1* also resulted in diminished biofilm development and ROS-mediated killing by myeloid phagocytes, thus, elucidating the role of Hog1 as an important regulator of *C. auris* skin colonization, biofilm formation, and systemic infection.

Unlike most *Candida* species, *C. auris* displays unique skin tropism and can persist on the human skin for a long time. This attribute is thought to be an important factor for *C. auris* outbreaks in healthcare settings (17), which has also been responsible for life-threatening systemic infections in susceptible patients. However, the skin adhesion or colonization mechanisms, as well as systemic virulence attributes of *C. auris* have remained poorly understood. The genome of *C. auris* is reported to contain adhesins belonging to *ALS* and the *IFF*/*HYR* family (21, 23). However, unlike *C. albicans*, which contains at least eleven Als adhesins, thus far only three *ALS* genes have been identified in *C. auris* (21, 23). Furthermore, the *C. auris* genome codes for several species-specific cell surface glycoproteins that may facilitate *C. auris* adhesion to different substrates (21, 23). Notably, there are clade specific differences in adhesin expression between the *C. auris* isolates (21). In this context, a *C. auris* specific adhesin, Scf1 and the Iff family member B9J08_004109 (IFF4109), were recently identified for their roles in surface colonization, adhesion, and virulence (23). We observed transcript levels for *IFF4109* to be significantly decreased in *hog1*Δ while the *SCF1* transcripts were not affected in the *hog1*Δ compared to the wildtype parental strain. Both *SCF1* and *IFF4109* are functionally redundant for biofilm formation (23), thus the transcriptional downregulation *IFF4109* alone does not explain the defective biofilm phenotype of the *hog1*Δ mutant. The *C. auris* cAMP/PKA signaling pathway has been shown to be important for biofilm formation (61, 62). However, in the *hog1*Δ we observe a modest increase in the transcript of B9J08_004030 (*TPK1*) (Table S1). In future studies, it would be of interest to determine genetic interaction(s), if any, between the *C. auris* Hog1 and PKA pathway. Furthermore, it is likely that Hog1 regulates additional, yet-to-be-identified, molecules for biofilm formation. Future studies are warranted where careful examination of the Hog1-regulated molecules will provide such insights.

In *C. albicans*, Hog1 MAPK is a major regulator of fungal stress response, morphogenesis, and virulence (33–35, 37, 63). The *C. auris* Hog1 has previously been reported to be important for antifungal drug resistance, stress response, and was also shown to be important for virulence in a *Caenorhabditis elegans* model of infection (40, 41). In addition to these previous findings, our genome-wide transcriptional profiling of logarithmically growing *hog1*Δ cells suggested that Hog1 modulates transcription of genes with putative functions in cell wall organization, adhesion, ROS-mediated phagocytic killing, and fungal pathogenesis, evident by the enrichment of distinct GO terms in the *hog1*Δ mutant compared to the wildtype parent. These data imply specific functions for Hog1 in different processes, including cell wall organization, cell adhesion, and pathogenesis. Indeed, the transcriptional data correlates well with *hog1*Δ phenotypes such as decreased biofilm formation, as described above. Additionally, the Hog1 functions in cell-wall organization are supported by our observations of enhanced β-glucan exposure and decreased total chitin levels in *hog1*Δ cells. It is likely that the reduced expression of β-1,3 glucanase B9J08_001418 (*BGL2*), and B9J08_003251 (*XOG1*), and chitin synthase B9J08_003879 (*CHS2*) contributes to increased β-glucan exposure and reduced chitin content in *hog1*Δ. Such cell wall changes can be highly functionally relevant, as have been shown previously using cell-wall mannosylation mutants (*C. auris* lacking *PMR1, PMT1, VAN1*), and distinct clinical *C. auris* isolates harboring unique cell-wall mannan structures (64–66). The increased β-glucan in the cell-wall mannosylation mutant strains has been shown to enhance neutrophil recruitment in a zebrafish infection model (65), and was associated with enhanced fungal killing by neutrophils and macrophages (64, 65). Interestingly, the loss of *HOG1* resulted in the increased β-glucan without impacting the total mannan levels, which highlights the hitherto unidentified mechanism by which Hog1 may regulate β-glucan exposure. Furthermore, the *hog1*Δ skin colonization, intradermal, and systemic infection did not result in enhanced neutrophil (or other phagocyte subset) accumulation in the skin or the kidneys despite the enhanced β-glucan exposure. These highlight the context-specific mechanisms that underly phagocyte accumulation in diverse *C. auris* infection settings. In contrast to the dispensable role in phagocyte accumulation, the enhanced β-glucan exposure in the *hog1*Δ mutant promoted the effector functions of neutrophils and macrophages and led to an enhanced fungal killing by both the phagocyte subsets. Overall, our findings confirmed the essential immunostimulatory role that the cell surface β-glucan plays while interacting with the host and uncovered a function for Hog1 in its masking for promoting the fungal survival.

Taken together, data presented herein demonstrates the *C. auris* Hog1 MAPK as an important regulator of skin colonization and systemic infection. Importantly, the data from the skin and systemic infection murine models imply that inhibitors of Hog1 may have beneficial effects in clinical therapeutic settings. However, further experiments are required to identify additional factors functioning together with Hog1 in the regulation of skin colonization, intradermal persistence, and systemic virulence. This might also lead to the discovery of additional potential antifungal drug targets, which could be used to render *C. auris* clinical strains less adherent to the skin.

## Materials and Methods

### *Candida auris* strains, media, and growth conditions

The *Candida auris* strains 1184/P/15 and *hog1*Δ used in this study were described previously (41). All data in the current study was generated by using two independent isolates of *hog1*Δ. All *C. auris* strains were routinely grown on YPD medium (1% yeast extract, 2% peptone, and 2% glucose (BD Biosciences) at 30°C with or without shaking at 200 rpm. For solid medium, 2% Bacto agar (BD Biosciences) was added to YPD broth. Synthetic sweat medium was prepared as previously described (53).

### Transcriptional profiling using RNA sequencing

For transcriptional profiling experiments, the *C. auris* wildtype and *hog1*Δ strains were grown to logarithmic growth phage in YPD broth at 30°C. Total RNA was purified using the TRI reagent (Sigma). Quality of RNA was assessed on a Bioanalyzer using the RNA6000 Nanochip (Agilent), mRNA was enriched using oligo(dT) beads (NEB) and subsequently, double-stranded cDNA libraries were generated by using the NEBNext Ultra II RNA Library Prep Kit for Illumina (NEB) according to the manufacturer’s instructions. The qualified libraries were subjected to Illumina sequencing with a 150 bp paired-read length at the Azenta Life Sciences sequencing facility. Three biological replicates for each strain were sequenced.

Quality control of raw sequencing reads was done using FastQC v0.11.8 (67). TrueSeq (Illumina) adapters were trimmed using cutadapt v1.18 (https://cutadapt.readthedocs.io/en/stable/; settings: -q 30 -O 1) followed by read mapping onto the *C. auris* B8441 genome assembly (http://www.candidagenome.org/) using NextGenMap v0.5.5 (68). Optical read duplicates were removed using Picard tools (Broad Institute, https://broadinstitute.github.io/picard/, settings: MarkDuplicates REMOVE_DUPLICATES=true VALIDATION_STRINGENCY=LENIENT).

Read counting was done using HTseq (69) in the union mode and the genomic annotation from *C. auris* B8441 (settings: -f bam -t gene -i ID). The read counts were normalized, and DESeq2 differential gene expression analysis was done using DEBrowser V1.28.0 (https://debrowser.umassmed.edu/). The Wald test was used to generate *P* values and Log2 fold changes. Genes with adjusted *P* values < 0.05 and absolute log2 changes >0.585 (1.5-fold change) were called as differentially expressed genes for each comparison. Normalized read counts were used for principal component analysis (PCA) in DEBrowser (70).

For further downstream analysis, *C. auris* genes detected in the RNA-seq analysis were mapped to *C. albicans* orthologs from the Candida Genome Database (http://www.candidagenome.org). Gene ontology (GO) annotations were used the GO slim mapper tool (http://www.candidagenome.org/cgi-bin/GO/goTermMapper) and a Fisher’s exact t-test was performed using R program. Only GO categories with a *P* value < 0.05 were considered significant. The RNA-seq analysis results are presented Table S1. The raw RNA-seq data are deposited to the Gene Expression Omnibus (GEO) under the accession number GSE256470.

### Biofilm formation assays

Biofilm assays were performed as described previously with slight modifications (71). Briefly, the *C. auris hog1*Δ and wildtype parental strains were grown overnight in YPD broth at 37^0^C. The overnight cultures were washed twice with PBS. The washed cultures were then grown for initial adhesion in RPMI 1640 medium with shaking at 37^0^C in a 4-well chambered cover glass that was pretreated with fetal bovine serum (FBS). After 90 minutes of growth at 37^0^C the cultures were washed once with PBS to remove unadhered cells. Fresh RPMI 1640 medium was added to cover glass wells and the cultures were further grown overnight at 37^0^C. The cells were washed once with PBS and stained with calcofluor white prior to microscopy. Microscopy was performed on a Leica STELLARIS 8 microscope using a 405 nm laser line and a 25X 0.95 NA water immersion objective. The tiled image stacks were acquired with the voxel size of 0.303 μm x 0.303 μm x 0.765 μm in x, y, and z directions. Multiple tiles were stitched together, rendered in 3D, and color-coded for Z-stack thickness using the Leica LAS X software.

### Cell wall quantification assay

The cell wall components were quantified by using a flow cytometry-based approach as described previously (41). Briefly, the *C. auris* strains were grown to logarithmic growth phase in synthetic sweat medium at 37^0^C. The logarithmically growing cultures were washed and stained with concanavalin A-conjugated Texas Red, Fc-hDectin-1a, and calcofluor white to quantify the mannans, glucan, and chitin, respectively. These triple-stained cells were measured in a BD Fortessa flow cytometer (BD Biosciences) to quantify the amount of chitin, glucan, and mannan using the BV421 (violet 405 nm, 50-mW power), fluorescein isothiocyanate (FITC) (blue 488-nm wavelength, 50-mW power), and Texas Red (red 640-nm wavelength, 40-mW power) detectors, respectively. A minimum of 10,000 events were recorded for each sample, and the data were analyzed using FlowJo software (BD Biosciences). Unstained and single-stained samples served as controls, and the data were expressed as the mean fluorescence intensity (MFI) from three independent experiments.

### Confocal microscopy

The *C. auris* wildtype and *hog1*Δ strains were grown to logarithmic growth phase in synthetic sweat medium at 30^0^C. The cells were washed twice with PBS and stained with Fc-hDectin-1a (Invivogen) and Calcofluor white (Sigma) as described above for cell wall quantification assay. The stained cells were fixed and imaged using a Leica Stellaris 8 confocal microscope with 405 and 561 nm laser lines and a 63X 1.4 NA oil immersion objective. The image stacks were acquired with the voxel size of 0.019 μm x 0.019 μm x 0.5 μm in x, y, and z directions. The image stacks were exported to Imaris (Bitplane) to quantify the fluorescence intensity. The fungal structures were segmented by creating surfaces around the calcofluor white stained channel using the "Surface Creation" module, with an "intensity threshold" above 25, and excluding any structures with "number of voxels" below 50. For the created surfaces around the calcofluor white positive fungal structures, the average "mean fluorescence intensity" was extracted for 10 randomly chosen regions of interest and were exported to Microsoft Excel.

### Macrophage/neutrophil killing assay

Primary cultures of bone marrow-derived macrophages (BMDMs) were isolated and cultivated exactly as described before (52). Neutrophils were isolated using a mouse neutrophil isolation kit (Miltenyi Biotec) according to manufacturer instructions. Survival of *C*. *auris* in BMDMs and neutrophils was quantified as described previously (52) using an MOI of 5:1 (fungi to macrophages or neutrophils). Fungal cells were harvested 2.5 h post infection and viability quantified by colony forming units (CFU)-counting of samples on YPD agar plates. Survival was calculated as percentage of viable CFUs after 48 h infection by comparing with uninfected *Candida* strains.

### Amplex^TM^ Red fluorescence assay

Hydrogen Peroxide/Peroxidase Assay Kit (ThermoFisher Scientific) was used to assess the H_2_O_2_ production as described previously (72). Briefly, 1x10^5^ macrophages or neutrophils resuspended in Krebs-Ringer phosphate glucose (KRPG) buffer (145 nM NaCl, 5.7 mM Sodium phosphate, 4.86 mM KCl, 0.54 mM CaCl_2_, 1.22 mM MgSO_4_, and 5.5 mM glucose), seeded in 96-well black plates along with 0.2 U/mL horseradish peroxidase and 50 μM Amplex^TM^ Red. Logarithmically growing *C. auris* strains were washed and resuspended in KRPG buffer. MOI 10:1 (fungi to macrophages/neutrophils) Candida cells were added to the assay plate. The appearance of the fluorescent resorufin was excited at 530 nm and emission was recorded at 590 nm. The fluorescence was measured every 2.5 minutes for the period of 2h at 37 °C using Tecan Infinite^®^ 200 PRO plate reader. Using H_2_O_2_ serial dilutions, a standard curve of resorufin fluorescence vs. H_2_O_2_ concentrations was constructed and was used to interpolate the H_2_O_2_ generated by macrophages/neutrophils after 2h.

### Murine models of epicutaneous, intradermal, systemic infections, and fungal burden determination

All animal experiments were performed according to the guidelines approved by the Center of Discovery and Innovation (CDI) and Purdue University’s Institutional Animal Care and Use Committee (IACUC). For all animal experiments, 8- to 12-week-old C57BL/6 (Jackson Laboratory) wildtype female mice were used. The infections were performed as described previously (58–60), with minor modifications. Briefly, in the case of skin infections, two to three days prior to infection, a depilatory lotion (Nair) was applied to mouse dorsal skin to remove hair from the anesthetized mice. Prior to infection with *C. auris,* the mice were anesthetized using xylazine/ketamine cocktail. The mice were infected with 5x10^7^ *C. auris* CFUs by topical application (50 μl) to the skin (epicutaneous infection) as well as 1x10^7^ *C. auris* CFUs via intradermal injection (100 μl). A separate group of mice were infected with *C. auris* retro-orbitally. At 72h post infection, the mice were sacrificed to quantify fungal burdens from infected skin and kidneys. The fungal burdens were quantified by plating the homogenates on YPD agar plates containing penicillin and streptomycin, and enumerating the CFUs after 48h of growth at 30°C.

### Single-cell preparation for immunophenotyping

Single-cell suspension from mouse skin and kidneys were prepared as described previously (59, 72, 73). Briefly, the skin tissue was collected and placed in 1 mL cold sterile digestion media (RPMI-1640 with 0.25 mg/mL Liberase TL Millipore Sigma) and 1 µg/mL DNase (Millipore Sigma). Skin tissue was then minced, 1.5 mL of additional digestion medium was added, and the minced tissue samples were incubated in a CO_2_ incubator for 100 minutes at 37 °C. Subsequently, 0.5 mL of 0.25% trypsin-1-mM EDTA was added to each sample and the incubation continued for additional 10 minutes, after which 2 mL of 1× PBS containing 5% FBS was added to each sample. For further tissue dissociation, the tissue pieces were taken in a 10-mL syringe, and were passed through the syringe 10 times, and were strained through a 40 µm cell strainer. The cell suspensions were washed twice and stained for flow cytometry.

For kidney single cell preparation, the kidneys were isolated and minced into <1 mm pieces. The kidney pieces were then transferred into a 50 mL conical tube containing 10 mL digestion medium (RPMI + 25 mM HEPES + 0.1 mg/mL DNase + 0.2 mg/mL Liberase TL) and incubated in a 37 °C shaking water-bath for 20 minutes. The digestion was terminated by placing the tubes on ice, and then ice-cold RPMI containing 10% FBS and 25 mM HEPES was added. The digested tissue pieces were then passed through a 70 μm cell strainer (Alkali Scientific) and treated with 5 mL ACK lysing buffer for 30 seconds (Quality Biologicals) for red blood cell lysis. After 30 seconds, 25 mL HBSS containing 2mM EDTA was added, and the suspension was then passed through a 40 μm cell strainer and resuspended in 40% Percoll (Millipore Sigma). The Percoll cell suspension was then overlaid onto 70% Percoll and centrifuged at 939 xg for 30 minutes at room temperature, with brake off. The kidney single-cell suspension was harvested from the interface, washed twice with FACS buffer (PBS containing 0.5% bovine serum albumin and 0.05% NaN_3_), and stained for flow cytometry.

### Flow cytometry-based immunophenotyping

The single cell suspension was stained with LIVE/DEAD Fixable Blue Dead Cell Stain (ThermoFisher Scientific) at 1:1000 dilution for 5 minutes on ice. Subsequently, to block non-specific antibody binding, the cells were incubated for 5 minutes on ice with rat anti-mouse CD16/CD32 (clone 2.4G2; BD Biosciences) at 1:100 dilution and 0.5% bovine serum albumin. For assessing the renal myeloid cells, the antibodies against specific cell surface antigens were used at 1:500 dilution. In the case of skin, the cells were stained with LIVE/DEAD Fixable Yellow Dead stain (ThermoFisher); after incubation with the rat anti-mouse CD16/CD32 Fc block, as above, the cell surface antigens were stained with the antibodies at 1:200 dilution. The following fluorochrome-coupled antibodies were used: Ly6G (clone 1A8, Biolegend), CD45 (clone 30-F11, Biolegend), CD11b (clone M1/70, Biolegend), CD3 (clone 145-2C11, Biolegend), CD19 (clone 1D3, Biolegend), CD11c (clone N418, ThermoFisher), Ly6C (clone AL-21, BD Biosciences), CD103 (clone 2E7, Biolegend), MHCII (clone M5/114.15.2, ThermoFisher), F4/80 (clone BM8, Biolegend), and NKp46 (clone 29A1.4, Biolegend). The cells were incubated with antibodies at 4 °C overnight, washed thrice with FACS buffer the next day, and passed through 35 μm cell strainer. The renal cells were then analyzed using the BD LSR Fortessa II (BD Biosciences), while dermal cells were analyzed using the Attune NxT Flow Cytometer (Invitrogen). The data were analyzed using FlowJo (BD Biosciences). The dermal phagocyte populations were determined as described before (59), while the gating strategy to identify the different renal myeloid phagocytes is provided in Figure S3.

### Histopathology

At 72 hours post-infection, skin and the right kidneys were harvested and placed in 10% buffered formalin overnight for fixation, followed by replacement with 70% ethanol. Paraffin-embedding, sectioning, and staining with Periodic acid-Schiff stain were carried out at Histoserve Inc (Gaithersburg, MD). The stained slides were scanned using Leica Aperio LV1 slide scanner at 40X. The scanned slides were opened in Leica ImageScope, from which the images were exported; FIJI (74) was then used to add scale-bars and the overlays, as depicted in the figure 4.

### Statistical analysis

Statistics were computed using GraphPad PRISM (version 10.0.3) and R in the case of RNA-Seq analysis. For assessing differences between groups, unpaired *t* test, Mann-Whitney *U* test, and ordinary one-way ANOVA with Tukey’s multiple comparisons test were used, as appropriate.

## Acknowledgements

This work was supported by grants from the National Institutes of Health (NIH) to NC (R21AI174118), JVD (R00AI141622), and ST (R01AI177604). This work is also supported in part by a grant from the New Jersey Health Foundation (NJHF) PC186-24 to NC. We thank the support from the Center for Discovery and Innovation (CDI) flow cytometry core for cell wall analysis and the CDI RAF for mice experiments. We thank all laboratory members for helpful discussions.

## Supplemental Material Legends

**Table S1.** List of differentially expressed genes in the *hog1*Δ mutant strain

## Supplementary figure legends

**Fig. S1:**
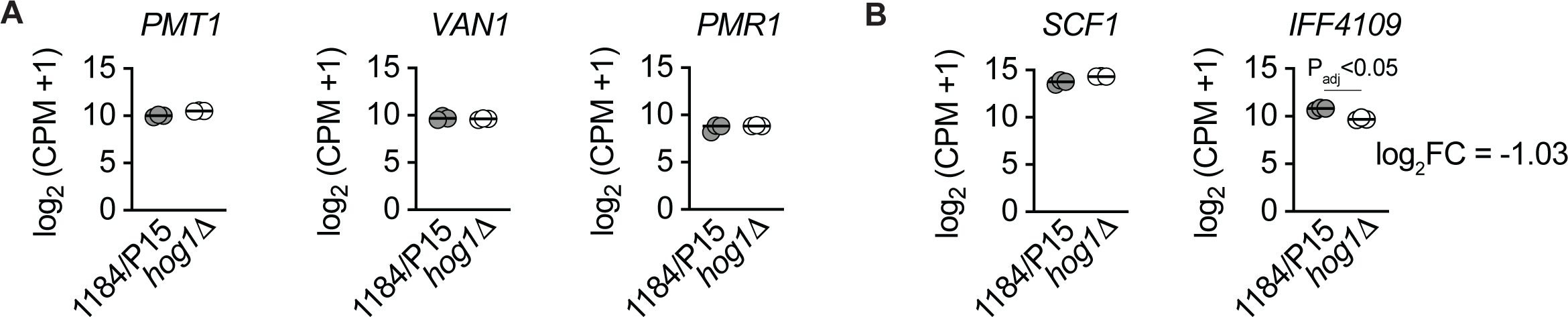
Transcript levels of genes involved in mannosylation and adhesion. (A-B) Gene expression profiles of *C. auris* isolates grown in YPD were determined using RNA-seq of three biological replicates. Log-transformed normalized counts (CPM+1) for indicated gene depicts the variation of *HOG1* expression in parent and *hog1*Δ cells. Dots represent individual replicate and horizontal line indicates mean. Approximately 2-fold (Log2FC: -1.03) reduction in the expression of *IFF4109* in *hog1*Δ cells compared to parent, indicated on the graph.

**Fig. S2:**
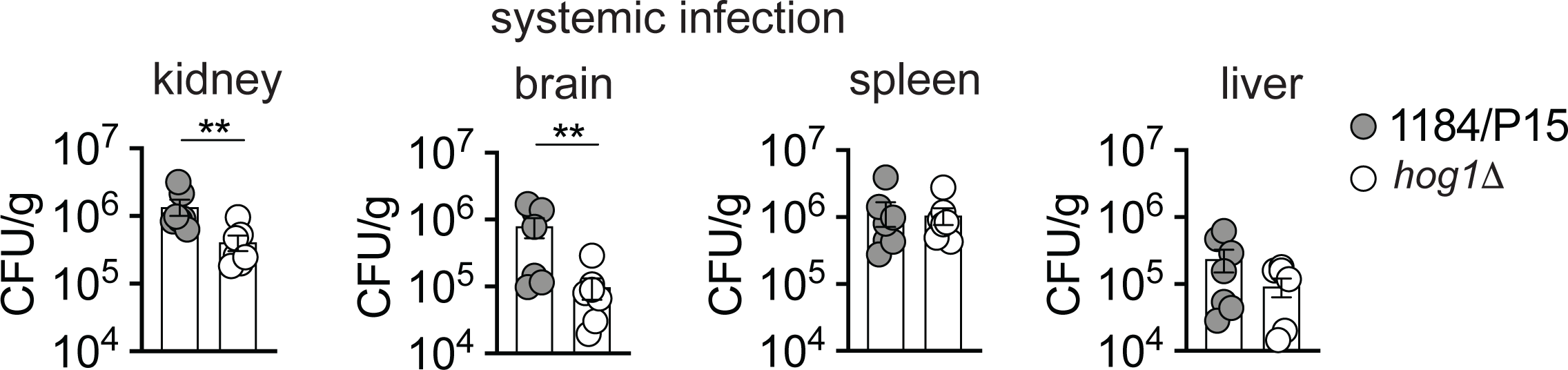
Genetic removal of *C. auris* Hog1 MAPK impairs murine systemic infection. (A-B) Fungal burden, at 72 hours post-infection for the indicated organs, after systemic infection of the immunocompetent C57/BL6 mice with 5 x 10^7^ *C. auris*. Data represents the mean of 6 individual mice. Dots represent individual mice. Error bars represent mean ± SEM; \*\**P* < 0.001, by unpaired Mann-Whitney *U* test.

**Fig. S3:**
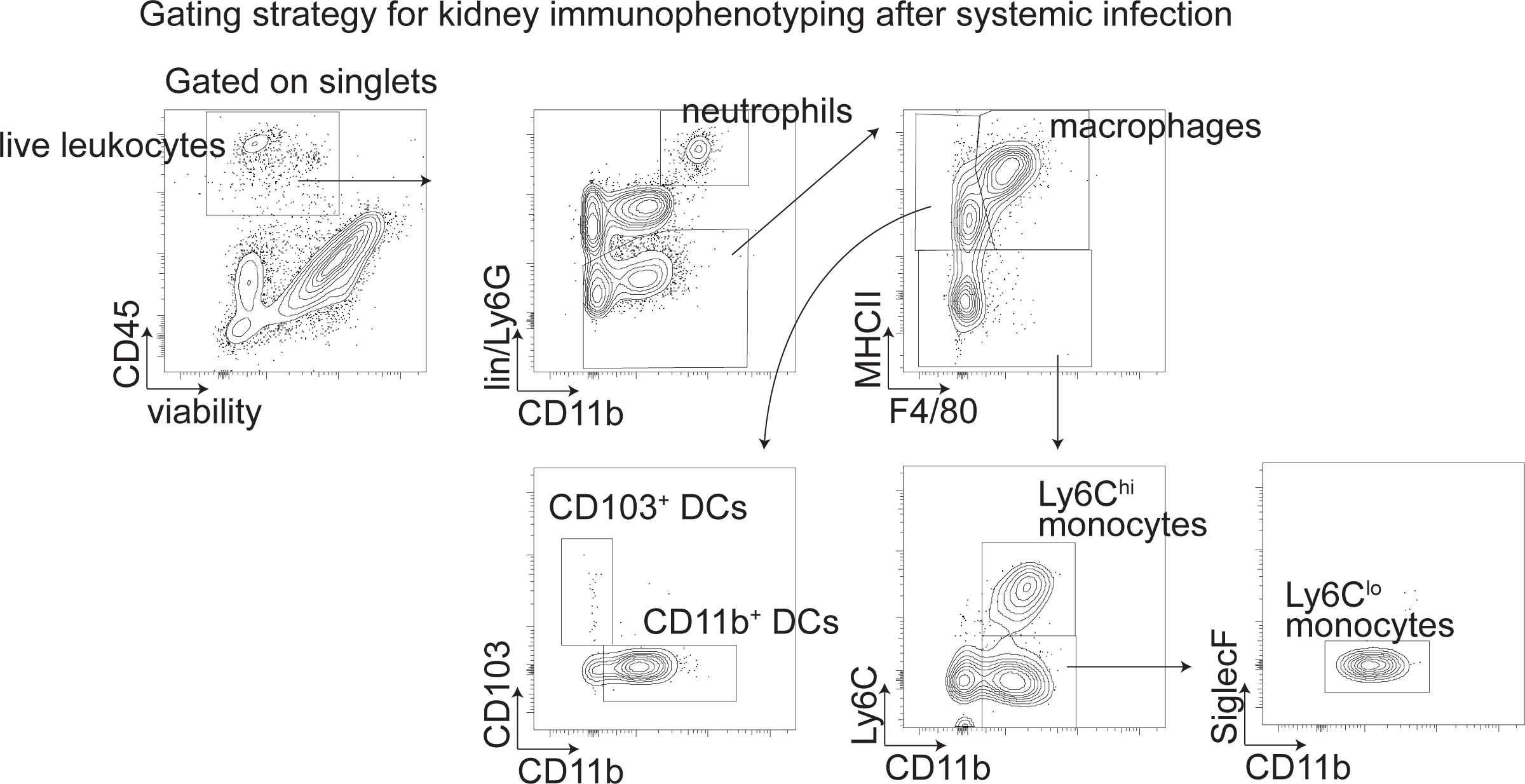
Immunophenotyping of *C. auris* infected kidneys. Gating strategy for the flow-cytometry based myeloid cell subsets in the mouse kidney.

**Fig. S4:**
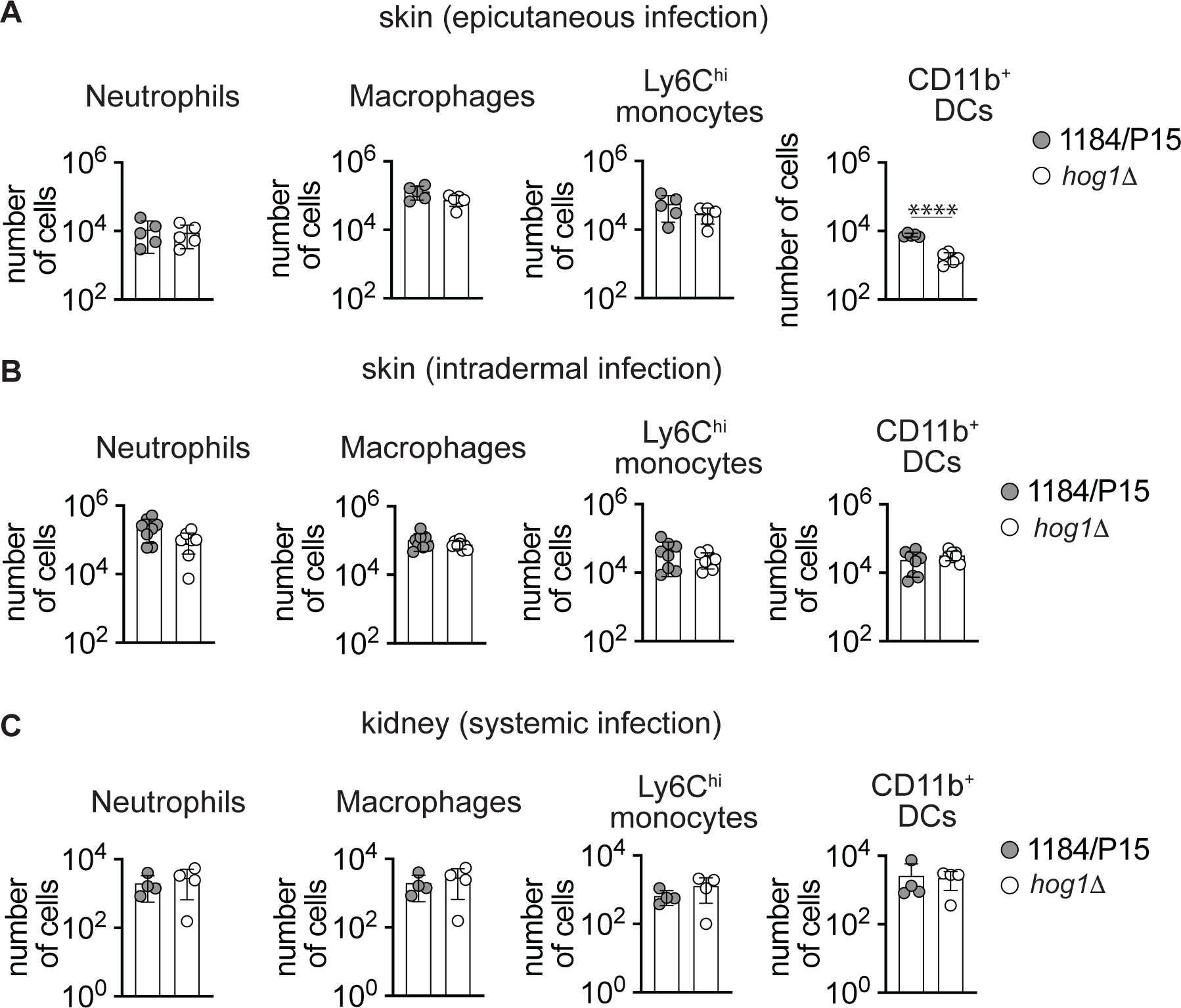
Myeloid phagocyte accumulation of *C. auris* infected murine skin and kidneys. The number of the indicated myeloid phagocyte subset after epicutaneous (A), intradermal (B) and systemic (C) infection, at 72 h post-infection, with the indicated strains of *C. auris*. Data represent the mean of 4-8 individual mice. Each dot represents an individual mouse. Error bar represents mean ± SD; ****p < 0.0001 by unpaired *t* test.

